# Fast barcode calling based on *k*-mer distances

**DOI:** 10.1101/2025.05.12.653416

**Authors:** Riko Corwin Uphoff, Steffen Schüler, Ivo Grosse, Matthias Müller-Hannemann

## Abstract

DNA barcodes, which are short DNA strings, are regularly used as tags in pooled sequencing experiments to enable the identification of reads originating from the same sample. A crucial task in the subsequent analysis of pooled sequences is barcode calling, where one must identify the corresponding barcode for each read. This task is computationally challenging when the probability of synthesis and sequencing errors is high, like in photolithographic microarray synthesis. Identifying the most similar barcode for each read is a theoretically attractive solution for barcode calling. However, an all-to-all exact similarity calculation is practically infeasible for applications with millions of barcodes and billions of reads. Hence, several computational approaches for barcode calling have been proposed, but the challenge of developing an efficient and precise computational approach remains. Here, we propose a simple, yet highly effective new barcode calling approach that uses a filtering technique based on precomputed *k*-mer lists. We find that this approach has a slightly higher accuracy than the state-of-the-art approach, is more than 500 times faster than that, and allows barcode calling for one million barcodes and one billion reads per day on a server GPU.

## 1 Introduction

DNA barcodes are used for identifying individual biomolecules in large populations. Applications include studying gene expression at the single-cell level [11], lineage tracing and screening [10], spatial transcriptomics [14, 20], DNA data storage [3], and the exploration of developmental trajectories and cancer progression [4, 19].

In a typical experiment, hundreds of millions of reads are sequenced, each containing one barcode modified by synthesis or sequencing errors that include substitutions, insertions, and deletions [17]. Correctly assigning the reads containing these modified barcodes to the barcodes they originate from is referred to as barcode identification or barcode calling [18]. Modern synthesis techniques such as photolithographic microarray synthesis allow the synthesis of millions of barcodes needed for spatial transcriptomics studies at a reasonable cost [13], but their error rates may exceed 20% per base [13], leading to the computational challenge of barcode calling for large barcode sets with high error rates. In a recent breakthrough result, Press [17] has shown that *random* barcodes of sufficient length are accurately decodable even for applications with such high error rates.

The central idea of many barcode calling approaches is to determine for each read a *most similar* barcode with respect to a given distance measure. However, the naive approach of identifying the most similar barcode for each read by computing the distance of each read and each barcode is prohibitively time-consuming, so several alternative approaches for barcode calling have been developed in the last decade: Zorita et al. [21] propose the calculation of all pairwise edit distances using prefix trees and several optimizations of the classical Needleman-Wunsch algorithm as well as clustering (“starcode”). Tambe and Pachter [18] develop the barcode-calling approach “sircel” and demonstrate its efficacy for small barcode sets and short barcodes. Press [17] proposes a filtering approach (triage) based on trimer similarities and thus accelerates barcode calling by orders of magnitude, thereby allowing barcode calling for one million barcodes and ten million reads per day on a server GPU. Yet, Press’ filtering approach is still computationally expensive as it requires a linear scan over the entire barcode set for each read, with a non-negligible workload per iteration.

Here, we propose a *k*-mer filtering approach that enables barcode calling for one million barcodes and one *billion* reads per day on an NVIDIA A100 server GPU.

This speed-up is accomplished by a simple auxiliary data structure that links given barcodes to their sub-sequences. Setting up this data structure requires only linear additional space and negligible running time in a preprocessing phase. We find by theoretical and experimental studies that the proposed *k*-mer filtering approach allows reducing the computational work by more than two orders of magnitude compared to Press [17]. An implementation of this barcode calling approach named “Qui*k*” can be accessed freely on the Github repository https://github.com/uni-halle/quik.

The rest of this paper is structured as follows: In Section 2, we provide a high-level description of the *k*-mer filtering approach and additional material used for the subsequent analyses and experiments. In Section 3, we present results on the accuracy of barcode calling using the *k*-mer filtering approach, results of a theoretical analysis of its running time as a function of *k*, and results of an empirical study of its running time. Finally, we conclude this article in Section 4. In the Supplementary Material we provide more detailed information on the simulation of reads, present further results, and give an analysis of Press [17].

## 2 Methods

In this section we present a high-level description of the barcode calling approach used in Qui*k*, details of the *k*-mer pseudo-distance used to filter candidate barcodes, several distance measures for quantifying sequence similarities, and combinations of different variants of the barcode calling approach, resulting in a two-step barcode calling scheme.

### 2.1 Barcode calling

The task of barcode calling is to identify for each read the barcode from which it originated. The Qui*k* approach assumes that all barcodes have a fixed length *ℓ*. Meanwhile, in a real experiment, reads are generally much longer DNA strings, each containing a barcode section of the same length *ℓ* at a well-defined position. To identify the barcode of a read, one can extract the barcode section of the longer DNA string. For simplicity, we will refer only to this extracted barcode section when referring to a specific read. On an abstract level, the Qui*k* approach can be divided into three phases.

- **Phase 0: Preprocessing**: For each possible *k*-mer *m* ∈ {*A, C, G, T*} ^*k*^ and each possible starting position *j* ∈ {0, …, *ℓ* − *k*} within a barcode, compile a list *L*(*m, j*) of *all barcodes* in which *m* occurs as a sub-sequence starting at position *j*.

While Phase 0 only needs to be executed once for a fixed barcode set, the other phases need to be executed *for all reads r*.

- **Phase 1: Barcode Filtering**: Evaluate for each barcode *b* some efficiently computable *pseudo-distance q*(*b, r*) (Section 2.2). Compile a small candidate set *C* of at most *c* barcodes with *smallest* pseudo-distance to *r*.
- **Phase 2: Final assessment**: For each candidate barcode *b* in *C*, evaluate some distance *d*(*b, r*) (see Section 2.3). Return a candidate barcode with minimum distance to *r* if this distance is smaller than a given threshold *D*.

Our approach has some similarities to that of Press [17]. However, the main difference is the filtering step in Phase 1. While Press evaluates several criteria that are ultimately combined into a final barcode ranking, we construct a candidate set with much less computational effort. As we will show in Section 3, this accelerates the filtering step by a large factor without any loss of accuracy. In the remainder of this section, we describe some details of the Qui*k* barcode calling approach.

### 2.2 *k*-mer pseudo-distance

The original barcode will be included within the candidate set with a high probability if the pseudo-distance *q* and the exact distance *d* have a high correlation, so the challenge of devising a suitable filtering approach is to find a pseudo-distance that (i) can be computed sufficiently fast and (ii) is sufficiently similar to the distance *d*.

#### Algorithm 1 *k*-mer pseudo-distance

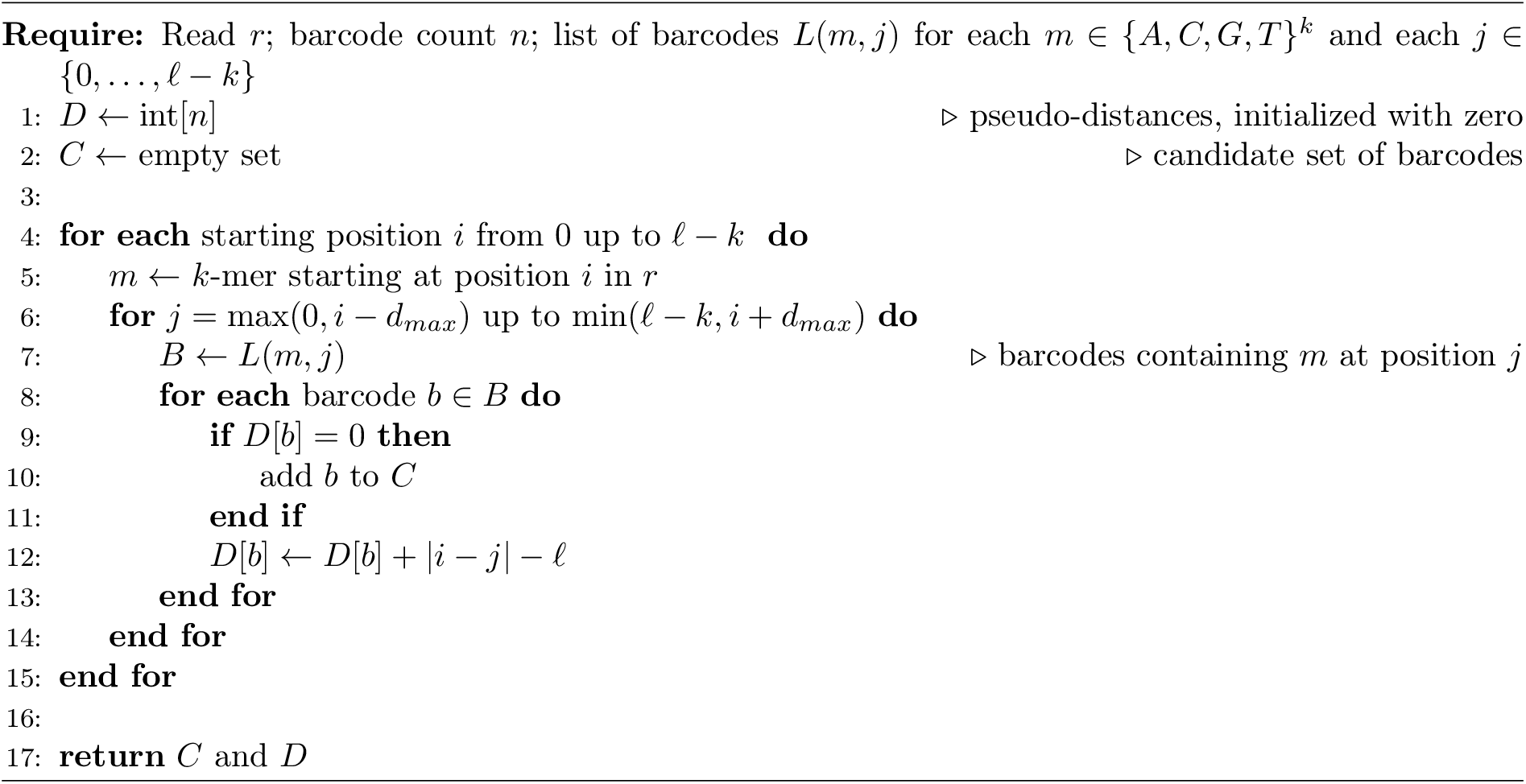

The central idea of our *k*-mer pseudo-distance *q*(*b, r*) is to consider each *k*-mer in the read *r* and to search for its nearest occurrence on the barcode *b*. The absolute difference between the starting positions of the *k*-mer in *r* and *b* quantifies how much this *k*-mer has been shifted away from its original position. We define the pseudo-distance as the sum of the absolute positional differences over all *k*-mers that occur in the read *r*. If some *k*-mer of *r* does not exist in *b* or is shifted away by more than *d*_*max*_ positions, we add *ℓ* as a penalty to the pseudo-distance instead.

The above description should be sufficient to explain the main idea and give some intuitions but is not enough to evaluate the pseudo-distances efficiently. To achieve this goal, we had to make some adjustments.

1. Instead of searching for the *closest* occurrence of a given *k*-mer in the barcode, we assume that each *k*-mer occurs *only once* within the threshold difference *d*_max_, and speed up the calculation by accounting for *all* occurrences of the *k*-mer within the threshold distance.
2. We make use of the precomputed lists *L*(*m, j*) of barcodes in which a *k*-mer *m* occurs at position *j*. This allows us to evaluate the pseudo-distances between a fixed read *r* and *all* barcodes in one run without having to search for *k*-mers on the barcodes.
3. We avoid explicitly adding the penalty term *ℓ* for *k*-mers that *do not* occur in the barcode or that are shifted by more than *d*_max_ positions. To do this, we subtract from each pseudo-distance *q*(*b, r*) a fixed value of *ℓ* for *each k*-mer in the read *r*, and thus *ℓ ·* (*ℓ* − *k* + 1) in total. This makes the penalty terms zero and so they no longer need to be treated explicitly. As a consequence, the pseudo-distance is always non-positive, with zero being the largest possible (and therefore worst) distance.

Alg. 1 implements these ideas and gives some additional details.

The algorithm constructs a candidate set *C* of barcodes that share at least one close *k*-mer with the read *r*. In the end, the array *D* contains the pseudo-distance of each barcode. The candidate set *C* can then be reduced to the *c* barcodes with the smallest pseudo-distance.

### 2.3 Distance measures

After compiling the candidate set, we compute a precise distance measure *d*(*b, r*) for each candidate barcode. Several distance measures have been proposed to quantify the distance between two given sequences, including the Hamming distance [6, 15], the Levenshtein distance [1, 8, 17], or the Sequence-Levenshtein distance [5]. While the Hamming distance is not suited to account for insertions or deletions (“indels”) in the read, the Levenshtein distance and the Sequence-Levenshtein distance do so. However, the Levenshtein distance requires the user to know the lengths of both sequences. As deletions and insertion can alter the length of the relevant segment in the read, the actual length of the substring corresponding to its barcode is unknown. To account for this, Buschmann and Bystrykh have proposed the Sequence-Levenshtein distance to allow truncation of both sequences at their ends at no cost [5]. Hawkins et al. have noted that the Sequence-Levenshtein distance is not a distance as it can violate the triangle inequality [9]. Still, they have used it for generating error-correcting barcodes, and we use it in this work as well.

Several ways to compute the (Sequence-)Levenshtein distance have been discussed. For barcodes of length *ℓ*, the classical dynamic programming algorithm computes the (Sequence-)Levenshtein distance in *O*(*ℓ*^2^). Under the Strong Exponential Time Hypothesis (SETH) this is optimal up to subpolynomial factors [2]. However, for a given threshold *D*, one can check in time *O*(*Dℓ*) whether two strings of length *ℓ* have a (Sequence-)Levensthein distance of at most *D* [12, 16]. As we are only interested in the (Sequence-)Levenshtein distance up to *D < ℓ*, we might also adapt the algorithm proposed by Zorita et al. [21] to accelerate the computation.

However, the sequence length *ℓ* is typically a small constant in practical situations (in our case *ℓ* = 34). In such situations, an optimized variant of the classical dynamic program is usually faster than the other, more advanced methods. Thus, we use the classical *O*(*ℓ*^2^) algorithm in our implementation.

### 2.4 Two-step barcode calling

We find from the results of Section 3 that increasing the length *k* of the subsequences trades better performance for a loss in precision and recall. Furthermore, one can raise the precision at the cost of diminished recall by decreasing the maximum accepted Sequence-Levenshtein distance *D*. Hence, we can tune Qui*k* to attain high throughput and high precision but low recall by choosing a large *k* and a small *D*.

We can exploit this observation to quickly identify the “easy” reads with few mutations and leave the rest of the missed reads for a slower but more accurate variant. We call the resulting scheme *two-step calling*. This calling scheme has two parameters for the threshold distances *D*_1_ and *D*_2_ and two parameters for the lengths of the subsequences *k*_1_ and *k*_2_ (with *k*_1_ *> k*_2_). In a first step, we run the barcode calling approach with the parameters *k*_1_ and *D*_1_ to quickly identify barcodes that are close to the associated read. For the reads that are not yet assigned, we run a second step with the parameters *k*_2_ and *D*_2_.

### 2.5 Parallelization strategies

To gain a high throughput of reads, it is essential to run a large number of calculations in parallel. Since Phases 1 and 2 are executed independently for each read, there are two obvious parallelization strategies for the Qui*k* calling scheme.

1. Press [17] describes how to parallelize the search for the most similar barcode for a fixed read. Following this idea, one would sequentially process one read after the other and do the calculations of Phase 1 and 2 in parallel.
2. We propose an orthogonal parallelization strategy in which we process the reads in parallel and, for each read, sequentially search for the most similar barcode. We thus launch a large number of long-running parallel threads, each responsible for one read, and each running Phases 1 and 2 sequentially.

While the first strategy is highly efficient when the number of reads is small, the resulting parallelization overhead makes it unattractive when the number of reads grows into the millions. In such situations, the second approach seems to be more promising. For this reason, we implemented the second strategy.

### 2.6 Error simulation

To assess the accuracy of barcode calling approaches, one needs labeled data sets where the original barcode is known for each read. Such data sets cannot be obtained experimentally, but they can be obtained by simulation [17], and we follow this approach here. As we employ a different error simulation strategy than Press [17], we give a brief overview of how we generate a read *r* from a barcode *b*.

We view the synthesis of a read as sending the barcode sequence through a noisy channel. Thus, we incrementally build up the read from the barcode with a certain probability of mutation *p* in each iteration. A mutation can be either a substitution (false interpretation of the signal), a deletion (missing a signal), or an insertion (false detection of a signal). To simplify the simulation, we assume that all three types of mutation are equally likely, each with a probability of *p/*3, and that the errors are independent across iterations. We also assume identical substitution probabilities for the twelve substitutions, identical insertion probabilities for the four insertions, and identical deletion probabilities for the four deletions. We note in passing that these assumptions are consistent with choosing the Levenshtein distance or the Sequence-Levenshtein distance for asserting the similarity between the read and the barcode. See Supplementary Material S1 for further detail on the simulation of reads.

While the assumptions typically do not hold in practice, the details can vary greatly between different experimental settings. Thus, we avoid making overly detailed assumptions about error distributions in order to maintain generalizability. One could also easily extend the procedure to more accurately capture the mutation probabilities of their experimental setting.

## 3 Results and discussion

In this section we present the results of several theoretical and empirical studies on our barcode calling approach. In particular, we discuss the accuracy of the *k*-mer pseudo-distance *q* and the *k*-mer filtering approach as well as a theoretical and empirical analysis of its running time.

### 3.1 Data set

In all of the following experiments and analyses, we used a fixed set of *n* = 1, 000, 000 barcodes of length *ℓ* = 34, drawn uniformly at random from the set of all barcodes of this length. To each of these barcodes, we applied the error model described in Section 2.6 with an error probability of *p* = 0.2 per base. In doing so, we generated a fixed set of 1, 000, 000 reads. The Supplement contains the results for the same experiments with error probabilities of 0.1 and 0.3.

### 3.2 Parameters

Both the accuracy and the running time of the Qui*k* barcode calling approach depend on the parameter choice, in particular, on the *k*-mer length *k* and the threshold distance *D*. In the following experiments, we tested several combinations of *k* ∈ {4, 5, 6} and *D* ∈ [0, 15]. For a clean and concise presentation, we fixed the parameters of the two-step variants to *k*_1_ = 7 and *D*_1_ = 10 and let the parameters *k*_2_ and *D*_2_ vary in the ranges mentioned above. In all experiments, we used a maximum shift size of *d*_max_ = 5 for the evaluation of the pseudo-distance (see Section 2.2). Finally, we used a candidate set of size *c* = 100 for all experiments unless explicitly stated otherwise.

### 3.3 Accuracy of the *k*-mer pseudo-distance *q*

In a first experiment, we studied the accuracy of the *k*-mer pseudo-distance *q* in comparison to the triage suggested by Press [17] in Phase 1. For this purpose, we applied each approach on our benchmark data set and determined a posteriori how often the original barcode occured within the candidate set of size *c* ∈ [1, 10^4^]. We call the relative frequency of this number the *hit rate* for size *c*. Thus, the larger the hit rate, the more often was the original barcode within the candidate set of size *c*. Consequently, a large hit rate is necessary for an accurate barcode calling. Fig. 1 shows the results of this experiment. We want to highlight some essential observations.

**Fig. 1:**
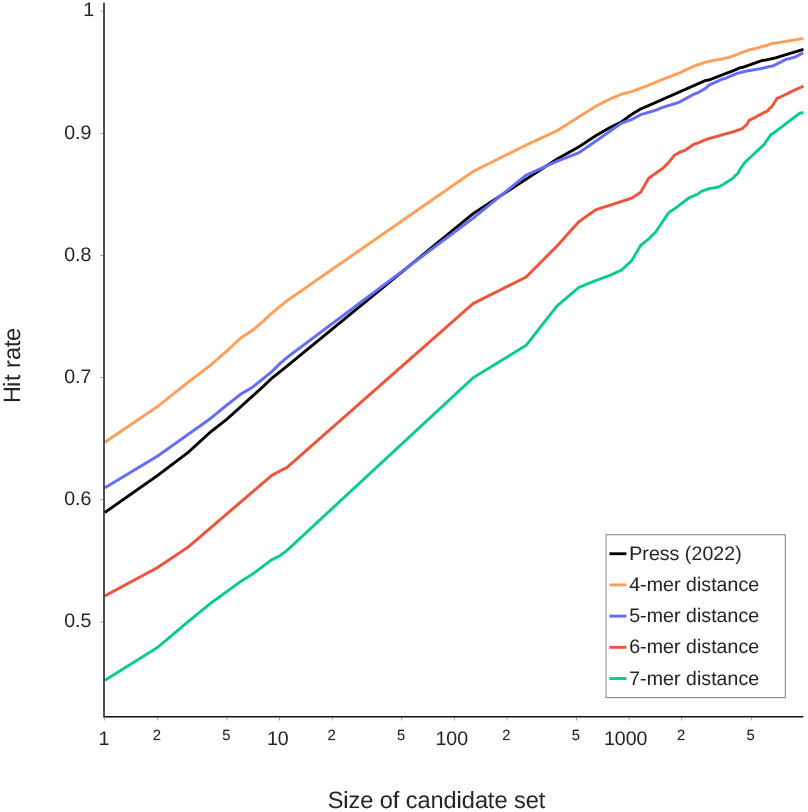
Relative frequency of the original barcode occurring in the candidate set of size *c* ≤ 10^4^. Note that the x-axis is scaled logarithmically.

- For all approaches, the hit rate grows monotonically and approximately linearly with log *c* for *c* ∈ [1, 10^3^].
- A candidate set of size *c* = 100 contains the correct barcode candidates in about 80% of the cases for *k* = 4.
- Increasing the value of *k* by one reduces the hit rate by a constant for a fixed size *c*.
- The hit rate for *k* = 5 is similar to that of the approach by Press [17].

We found qualitatively similar results by analogous studies with error probabilities of *p* = 0.1 and *p* = 0.3 (see Supplementary Material S2). We conclude that the pseudo-distance is well-suited for effectively pre-filtering the relevant barcodes. In particular, the hit rate is with *k* = 5 similarly high as with the triage rules suggested by Press.

### 3.4 Accuracy of the *k*-mer filtering approach

To study the accuracy of the *complete* barcode calling process, we apply the different approaches to the benchmark data set and calculate the *precision* defined as the ratio of correctly called reads and all accepted reads [17] and the *recall* defined as the ratio of accepted reads and all 1, 000, 000 reads [17]. Precision and recall highly depend on the choice of the exact distance measure and on the maximum acceptable distance parameter *D*. In this experiment, we calculated precision and recall for various values of *D* ∈ [0, 15] and for both the Levenshtein and Sequence-Levenshtein distance. Figs. 2a and 2b show the results.

**Fig. 2:**
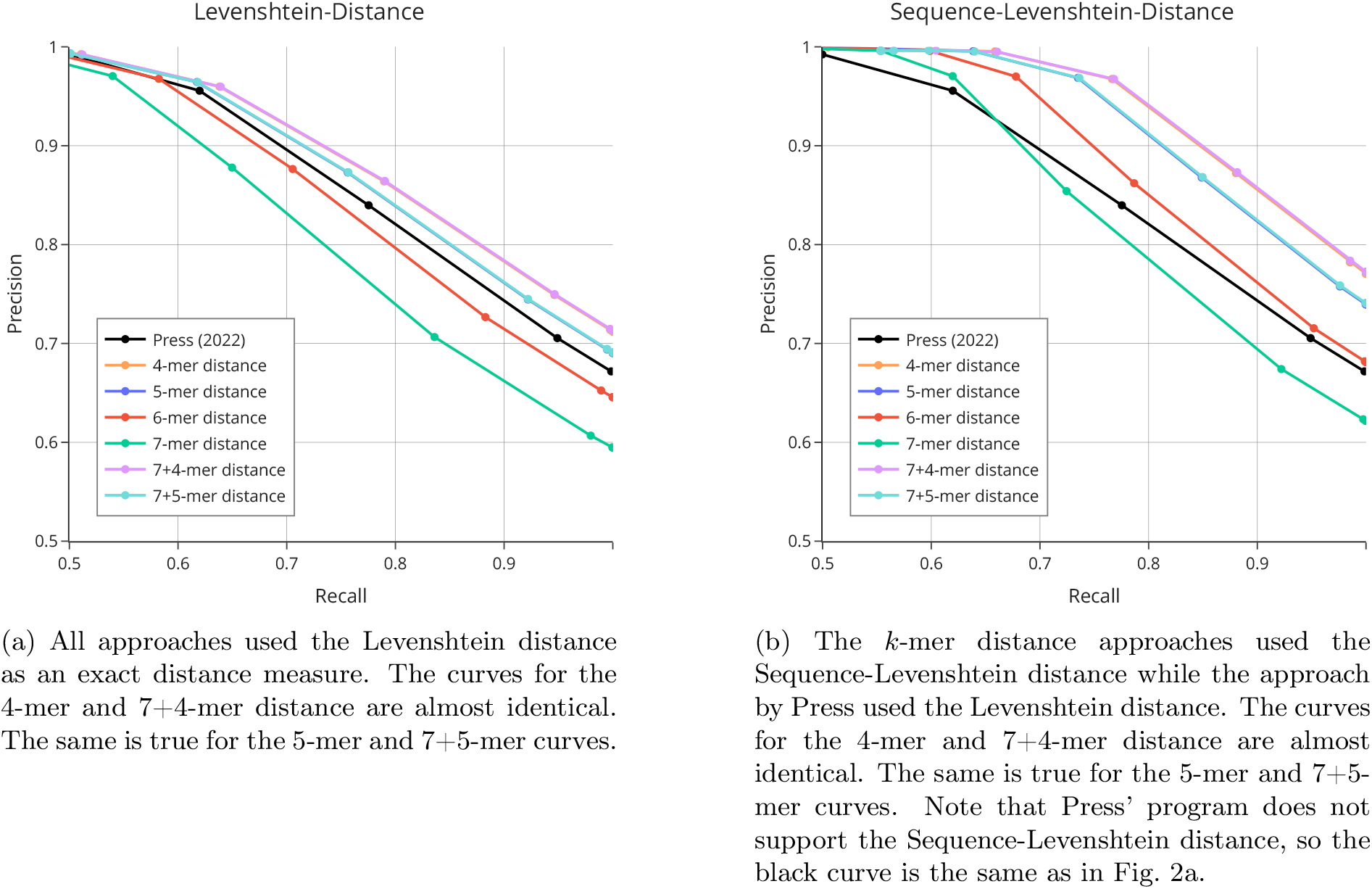
Precision-recall curves for different values of the maximum acceptable distance *D* to the read. Each integer value of *D* ∈ [0, 15] yields one point, forming a curve of 16 connected points for each approach.

The following observations are essential:

- At lower recall, all approaches yield a precision of ∼ 100%.
- The Sequence-Levenshtein distance gives higher precision and recall than the Levenshtein distance.
- Using the Sequence-Levenshtein distance, the *k*-mer distance approaches for *k* ∈ [4, 6] yield higher precision and recall than Press’ approach.
- We find that the precision and recall of the two-step variants is almost identical to the associated one-step approach.

Based on the results of this experiment, we used a maximum acceptable distance of *D* = 8 and the Sequence-Levenshtein distance in the following experiments. In doing so, we gain a barcode calling approach with similar or higher accuracy than Press.

### 3.5 Theoretical analysis of running time

In this subsection, we present a theoretical analysis of the expected running time required to calculate the pseudo-distances between a single read and *all* barcodes as described in Algorithm 1. For this purpose, we concentrate on the expected number of executions of Line 12 in Algorithm 1, which we call the number of *inner iterations*.

This analysis is based on the two simplifying assumptions that all *k*-mers of a sequence be independent and all *k*-mers be equally likely. In general, assuming independence will lead to an overestimation of the number of operations performed. The main result of this analysis is the following.

#### Theorem 1.

*Phase 1 is expected to take at most*

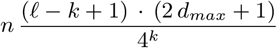

*inner iterations for computing the pseudo-distances for n barcodes of length ℓ*.

*Proof*. The probability that a random *k*-mer resembles a queried sub-sequence is about 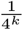, since there are 4^*k*^ different *k*-mers. Therefore, the list of barcodes containing a given *k*-mer at a fixed position has an average length of at most 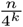. Hence, due to the assumed independence of the sub-sequences and since we consider at most 2*d*_*max*_ + 1 positions for each of the *ℓ* + *k* − 1 sub-sequences in the read, the number of inner iterations is at most

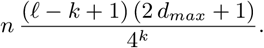

Theorem 1 implies that, under the assumption that *d*_*max*_ is a constant and that 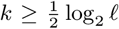 (a viable choice according to previous results), we obtain a time complexity of

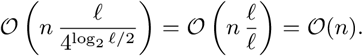

Theorem 1 also implies that increasing the parameter *k* by one reduces the expected number of iterations per read by a factor of four. Theorem 1 further implies that, for a parameter set of *ℓ* = 34, *k* = 4, and *d*_*max*_ = 5, we obtain on average at most 1.33*n* iterations per read. Using similar arguments, one can show that the approach by Press [17] takes 262*n* iterations per read (see Supplementary Material S3).

We conclude that the *k*-mer filtering step can *in theory* be executed about 262*/*1.33 ≈ 197 times faster than the triage rule filtering proposed by Press.

### 3.6 Empirical analysis of running time

To test whether the theoretical results from the previous section hold in practice, we next performed an *empirical* analysis of the number of inner iterations of Algorithm 1. To this end, we applied the approaches for several values of *k* to our benchmark data set and simply counted how often Line 12 is executed.

The following observations from Figs. 3 and 4 are essential:

**Fig. 3:**
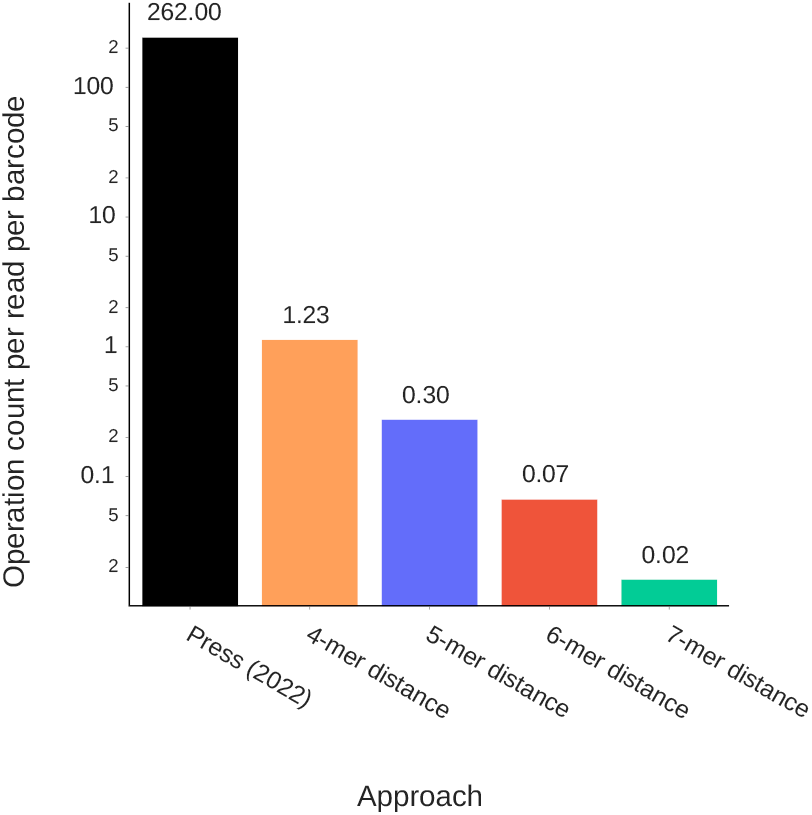
Normalized operation count. The ordinate shows the log-scaled mean numbers of inner iterations per read and barcode.

**Fig. 4:**
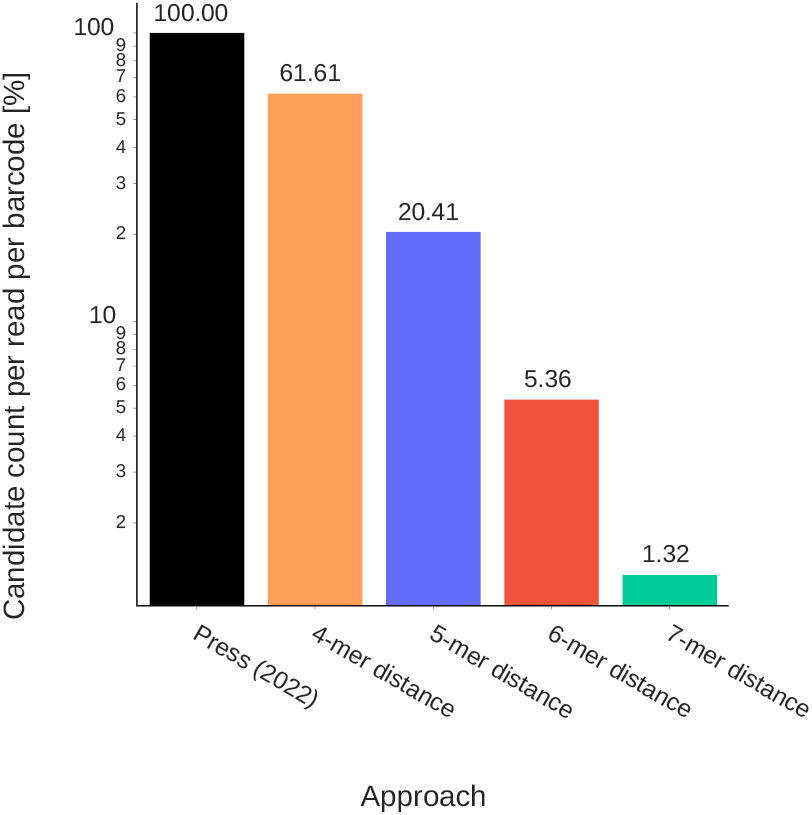
Fraction of barcodes that need to be scanned to find the best candidates. Note the log-scaled y-axis.

- The number of inner iterations decreases by a factor of 4 when increasing *k* by one. That is consistent with the theoretical analysis of Subsection 3.5.
- The computation of the 4-mer pseudo-distance requires about 1.23*n* inner iterations and thus about 262*/*1.23 ≈ 213 times fewer iterations than that of the approach by Press.
- When increasing *k* by one, the number of barcodes that share at least one *k*-mer with the read *r* reduces by a factor of about 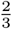.

For a given read, the approach by Press has to sort all barcodes by pseudo-distance to obtain the final candidate set. More accurately, the approach sorts five different candidate sets, each containing 100% of the barcodes, to combine them into the final pseudo-distance and sort all barcodes again. In contrast, the Qui*k* approach only once probes a much smaller fraction of all barcodes, namely those for which a *k*-mer of the read appears in the precomputed lists. In summary, our experiment confirms the previous theoretical analysis.

### 3.7 Empirical analysis of running time on GPUs

Next, we examined the *empirical* running time of the complete barcode calling approach as a function of *k* in comparison to Press’ approach. For this purpose, we implemented the *k*-mer filtering approach as a GPU program in CUDA and compared it to the GPU programm developed by Press. We applied each program to our benchmark data set and measured the running time of the calling process (Phase 0 to 2, but excluding the time for loading barcodes and reads from disk.) All computations have been executed on a Linux system with an AMD EPYC 7662 64-Core processor, 1 TB of main memory and an NVIDIA A100-SXM4 GPU having 6912 CUDA cores and 40 GB GPU memory.

Fig. 5a shows the results of this experiment.

**Fig. 5:**
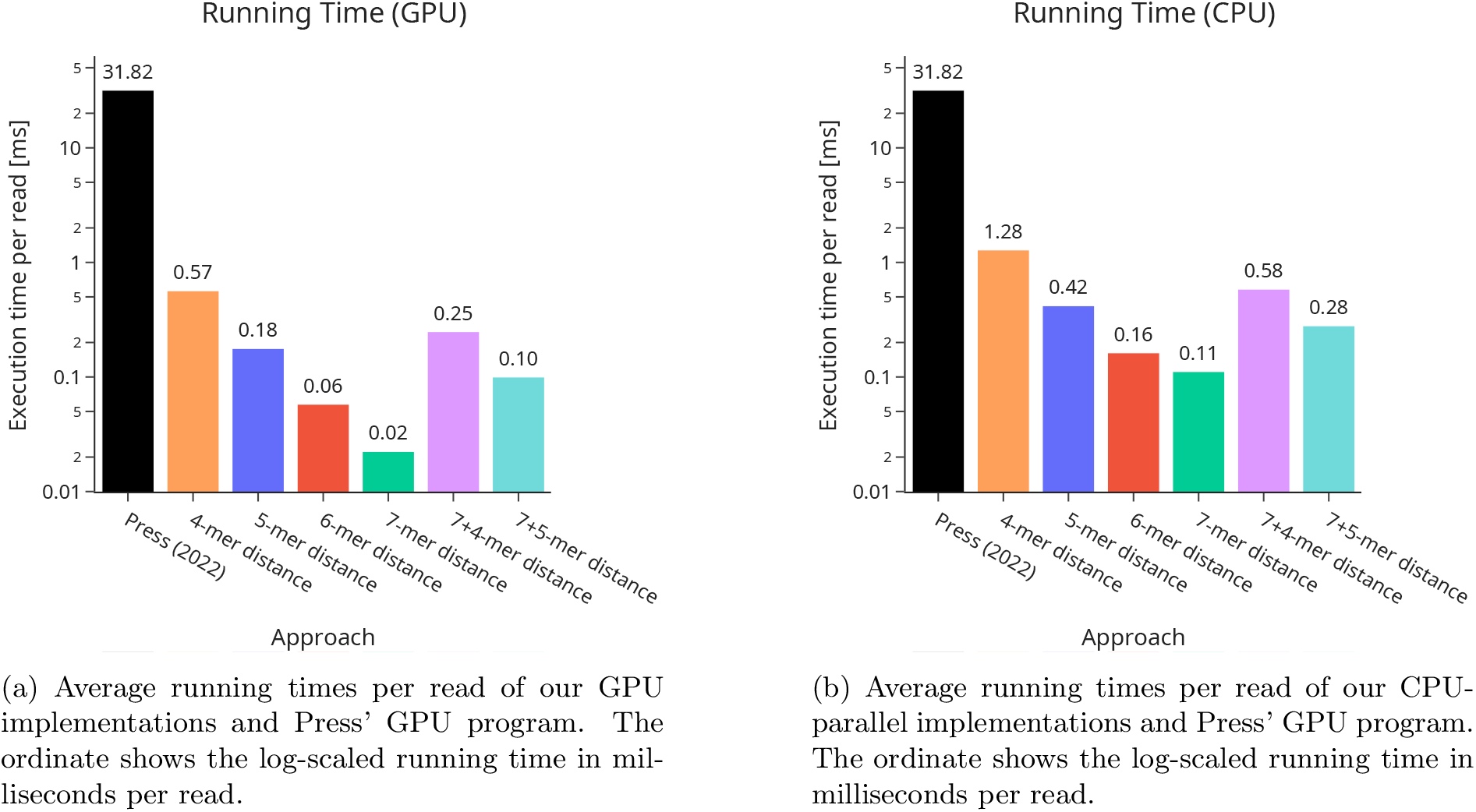
Comparison of running times.

- We observe that all *k*-mer approaches are much faster than Press’ program. In particular, the 6-mer calling approach (which has higher accuracy than Press’ program) is faster by a factor of 31.82*/*0.06 ≈ 530. This implies a theoretical throughput of about 24·60·60·1000*/*0.06 = 1.44·10^9^ reads per day.
- This large speed-up is induced by our efficient filtering step (Phase 1) and the different parallelization strategy. Our strategy also seems to be more GPU friendly as it utilizes 100% of the computing resources. Instead, Press’s program utilizes at best 33% of our GPU.
- We further observe that the two-stage filtering approaches (the two rightmost columns) can slightly reduce the running times. The combined 7+4-mer distance improves on the 4-mer distance, and the 7+5-mer distance improves on the 5-mer distance.

We conclude that the *k*-mer distance approach can not only be more accurate but also executed *much* faster than Press’ program on a server GPU.

### 3.8 Empirical analysis of running time on CPUs

Finally, we compared our GPU implementation of the *k*-mer filter approach with a classical CPU-parallel implementation using OpenMP. In contrast to ours, Press’ program still depends on the GPU.

We find from Fig. 5b that the running times of our CPU implementation are larger than the GPU times by a factor of less than three for the 6-mer distance. Our observations demonstrate that GPUs do not give an essential advantage to CPUs for the barcode calling process. This is good news for many applications, where GPUs are still rare.

## 4 Conclusions

The capability of performing barcode calling for billions of reads and millions of barcodes is essential in the modern life sciences in diverse areas including single cell and spatial transcriptomics, lineage tracing, or synthetic biology. This computational problem is particularly challenging when the error rate exceeds 10% or even 20% per nucleotide as in modern synthesis techniques such as photolithographic microarray synthesis. A recent breakthrough by Press [17] enables barcode calling in such cases for several million reads and one million barcodes per day per GPU. Here we have developed a method called Qui*k* based on *k*-mer filtering, which allows the user to tune the parameter *k* for choosing the trade-off between a desired level of accuracy and a low running time. This method is based on a pre-computed auxiliary data structure that allows an efficient filtering of relevant barcodes. In a theoretical analysis we have found that Qui*k*’s filtering step requires about 197 times fewer inner loop iterations than Press on average. In a first experimental analysis based on one million barcodes and simulated sets of reads with an error rate of 20% per nucleotide, we have found that the Qui*k* approach yields an accuracy slightly higher than that of Press and that reduces the running time on a GPU by more than two orders of magnitude, enabling the analysis of one billion reads per day per GPU. In a second experimental analysis, we found that the running time of a parallel CPU implementation using OpenMP is less than three times higher than that of the corresponding GPU implementation, but is still more than two orders of magnitude faster than that of Press. While large-scale analyses of single cell or spatial transcriptomics data with millions of barcodes would currently require a high-performance cluster for barcode calling on billions of reads with error rates above 10%, the increased throughput of the Qui*k* approach now enables these analyses on much cheaper hardware widely available in research labs worldwide.

## Acknowledgments

We thank Lothar Altschmied, Jan Grau, Rene Malsch, Paride Rizzo, and Antonia Schmidt for valuable discussions.

## Author contributions statement

R.C.U. and M.M.-H. developed the approach, R.C.U. and S.S. implemented the algorithms and performed the studies, and all authors analyzed the results and wrote the paper.

## Data availability

The CUDA implementation of the Qui*k* barcode calling approach, data files, and auxiliary scripts to reproduce our experiments are available on Github https://github.com/uni-halle/quik [7].

## Supplementary Material

### S1 Details on the simulation of reads

In this section we explain in detail how we generate a read *r* from a barcode *b*. As Figure 6 shows, we incrementally build up the read from the barcode with a certain mutation probability *p* in each iteration.

**Fig. 6:**
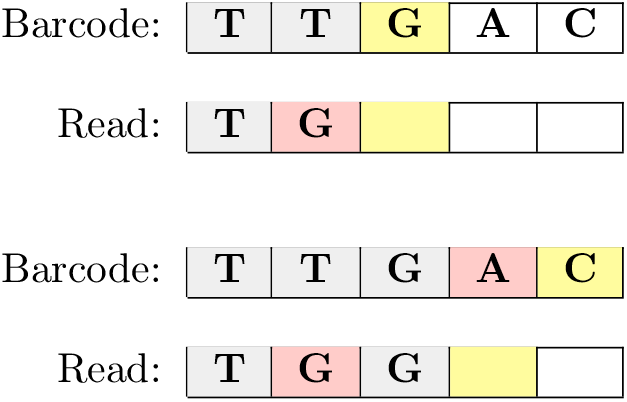
Iteratively building up the read. The positions marked yellow represent the index of the read/barcode considered in the current iteration. Red indicates that an error has occurred. Top: A substitution caused the base in the barcode to be replaced by a different base in the read (here: T is replaced by G). Bottom: A deletion caused the base in the barcode to be skipped (here: A is deleted).

If no mutation occurs, we progress to the next base of the barcode sequence and correctly appended to the read. Conversely, if a mutation occurs, it can be a substitution, a deletion, or an insertion. In the case of a substitution, we progress to the next base of the barcode sequence and append a different, uniformly random base to the read. In the case of a deletion, we proceed to the next base of the barcode, but the read remains the same. In the case of an insertion, the opposite happens with equal insertion probabilities for all bases.

However, if a deletion immediately follows an insertion (or vice versa), it would be impossible to distinguish it from a substitution. That would result in an incorrect assignment of one or even no error instead of two. Therefore, we do not allow insertions if the previous iteration resulted in a deletion, and vice versa. Finally, if the resulting read *r* is longer or shorter than the original barcode *b*, we truncate *r* to length *ℓ* or fill read *r* with bases drawn from a uniform distribution to achieve exactly length *ℓ*.

Algorithm 2 describes the resulting procedure we use to simulate erroneous reads. Reads are built up iteratively with fixed error probabilities in each iteration. In case of a deletion, the probability of insertion in the next iteration is set to 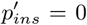, and the probability for a deletion becomes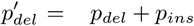. Insertions are treated analogously. In this way, we reduce the likelihood that a sequence of several successive mutations will erroneously lead to only one error or none at all. Thus, the output error rate will be closer to the desired input mutation probability.

Any viable error simulation model should approximate the expected error distribution of the underlying distance model. Since we use the Sequence-Levenshtein distance as a measure of similarity between sequences, one would aim for an average Sequence-Levenshtein distance of *p · ℓ* for the sequence length *ℓ* and a per-base error rate of *p*. To test the feasibility of the iterative error model, we empirically recorded the distribution of sequence-Levenshtein distances between uniformly random barcodes of length *ℓ* = 34 and the mutated reads with error probability *p* = 0.1.

As can be seen in Figure 7, the average Sequence-Levenshtein distance of the iterative error model deviates by less than 4% from the expected value of *p · ℓ* = 3.4. In contrast, the error simulation model described by Press [17] shows a 40% lower output error rate than expected. The low mutation rate in Press’ model could be explained by the way mutations are applied there. Because deletions and insertions occur after bases are substituted, the process tends to delete recently substituted positions, resulting in only one deletion in the output. In addition, many insertions in the final stage of the algorithm may cause mutated bases to exceed the desired length and thus end up truncated. However, the exact causalities are unclear.

**Fig. 7:**
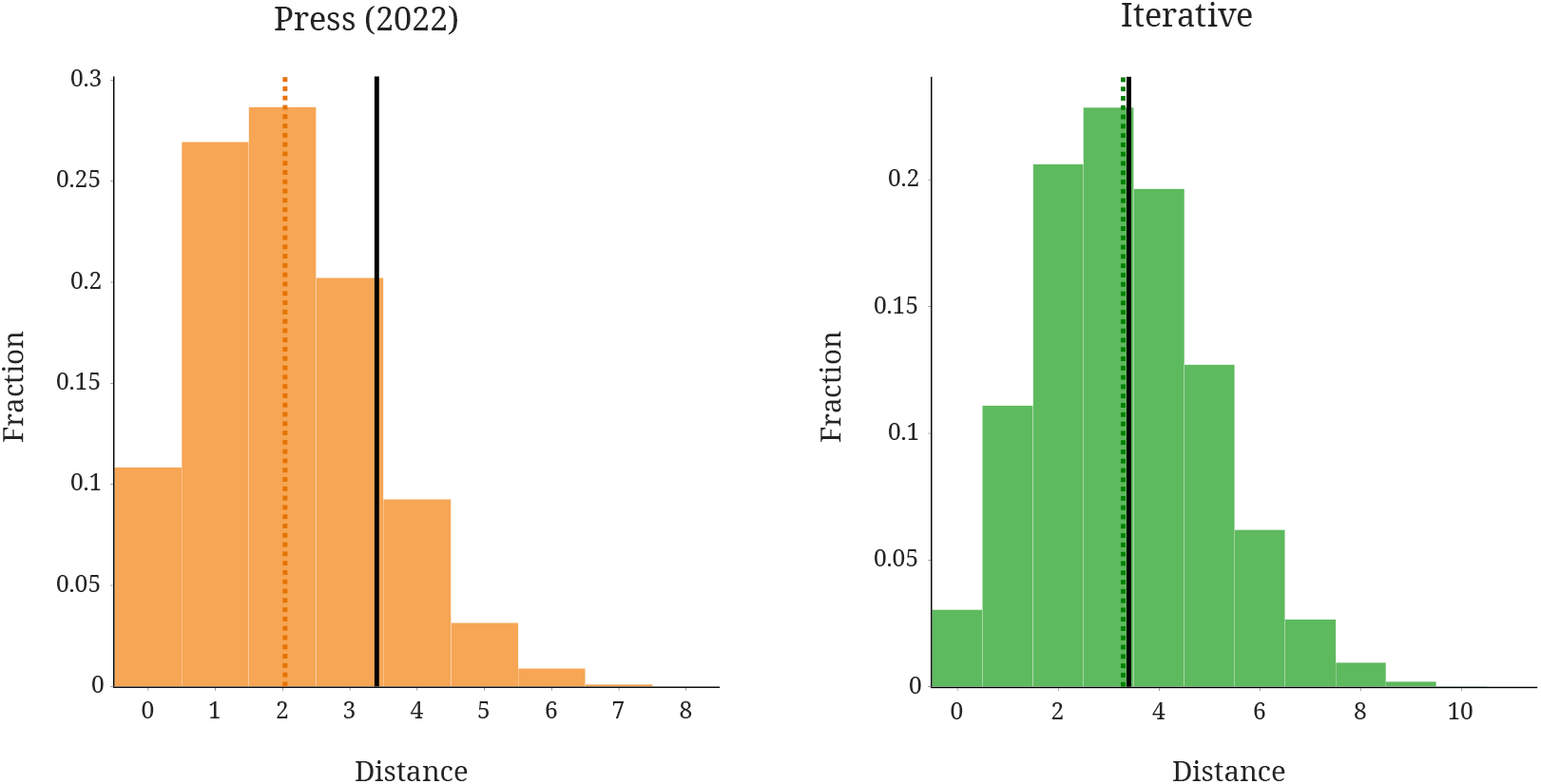
Relative frequencies of Sequence-Levenshtein distances observed when simulating the mutation of barcodes with length *ℓ* = 34 and error probability *p* = 0.1. Left: Results using Press’ model. Right: Distribution of the iterative procedure used in this study. The black line marks the expected mean distance of a perfect simulation model. The dashed lines represent the mean distances of the tested error simulation models.

#### Algorithm 2 Simulation of erroneous read

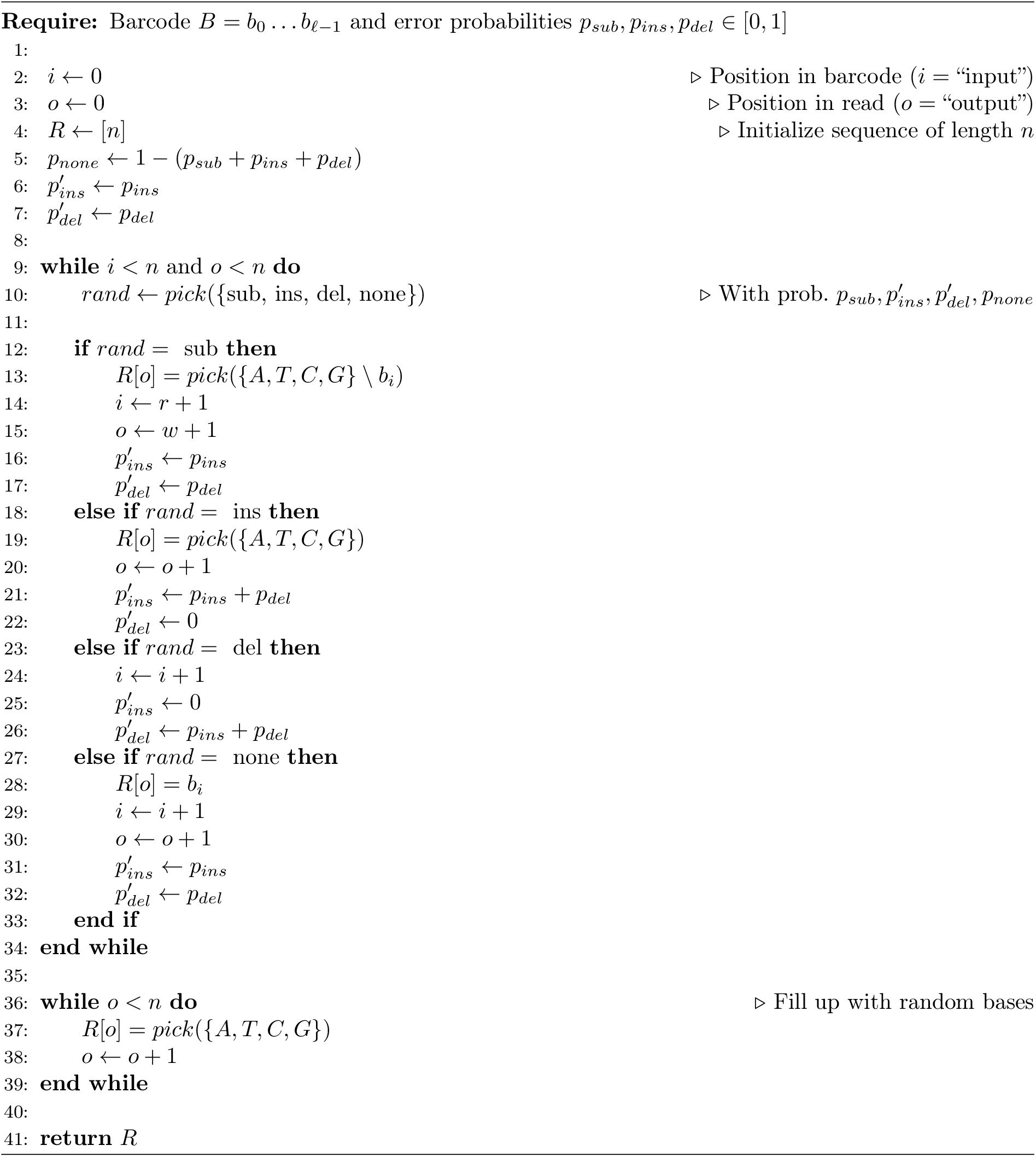

### S2 Supplementary results

The following section contains additional figures and experimental results. All of this additional material agrees with our general discussion in the main manuscript.

#### S2.1 Accuracy of the pseudo-distance

Figs. 8a and 8b show the results from the experiment described in Section 3.3 (Accuracy of the *k*-mer pseudo-distance *q*) when performed with a base error probability of 10 % and 30%. Although the absolute numbers differ, the relative order of the barcode calling algorithms stays the same.

**Fig. 8:**
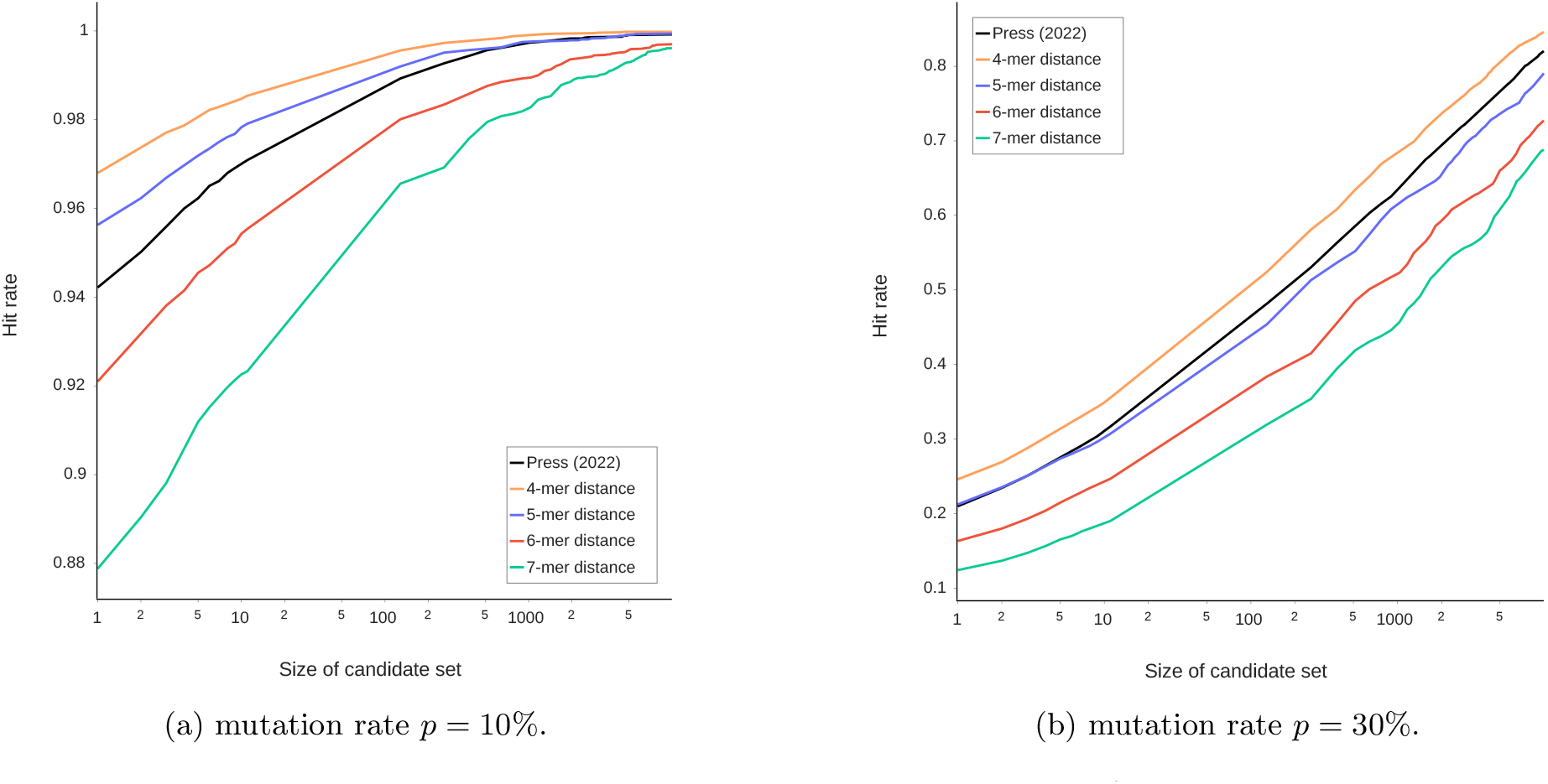
Hit rate at position *x* in the index for *n* = 10^6^ barcodes.

#### S2.2 Accuracy of the *k*-mer filtering approach

Figs. 9a and 10a show the results from the experiment described in Section 3.4 (Accuracy of the *k*-mer filtering approach) when performed with a base error probability of 10 % and 30% and the Levenshtein distance as an exact distance measure. In contrast, Figs. 9b and 10b show the results if we use the Sequence-Levenshtein distance. Again, the relative order is mostly unaffected by the change of the distance measure.

**Fig. 9:**
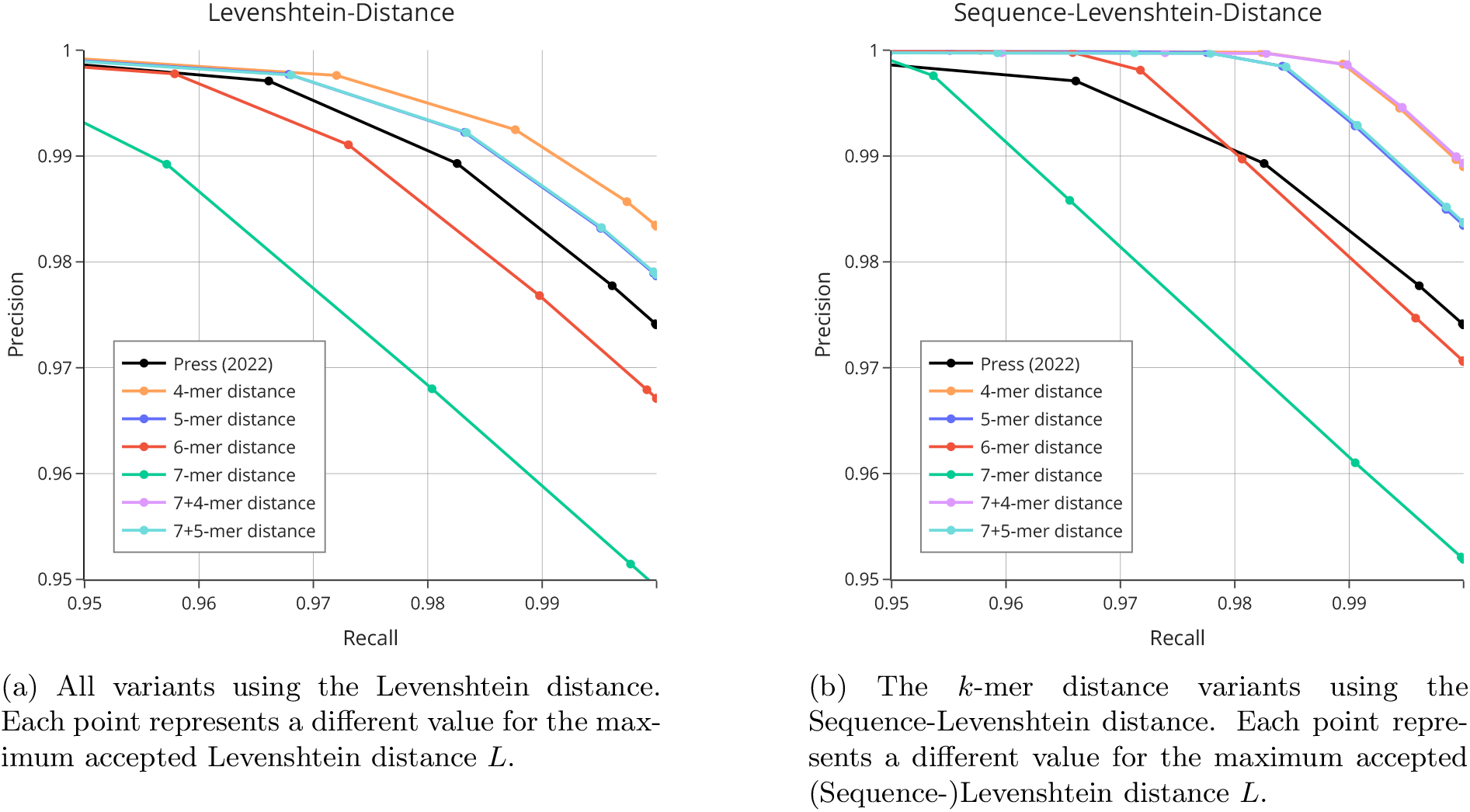
Precision-recall-curve for *n* = 10^6^ barcodes and reads and mutation rate *p* = 10%.

**Fig. 10:**
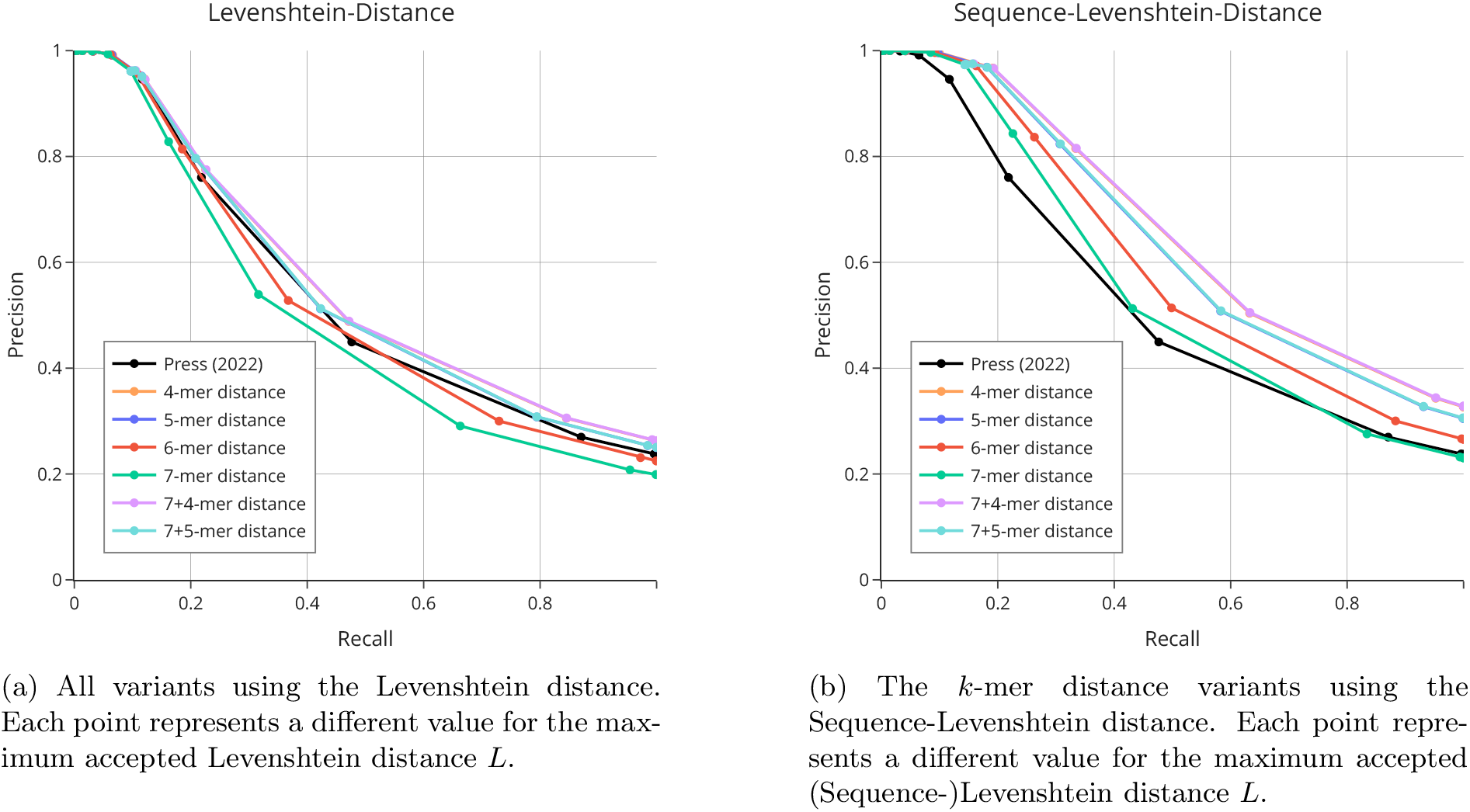
Precision-recall-curve for *n* = 10^6^ barcodes and reads and mutation rate *p* = 30%.

#### S2.3 Empirical analysis of runtime on GPUs

Figs. 11a to 12b show how the algorithm’s execution time on our benchmark GPU is affected by mutation rates of 10 % and 30 % and by the Sequence-Levenshtein distance. The results agree with our discussion in the main manuscript.

**Fig. 11:**
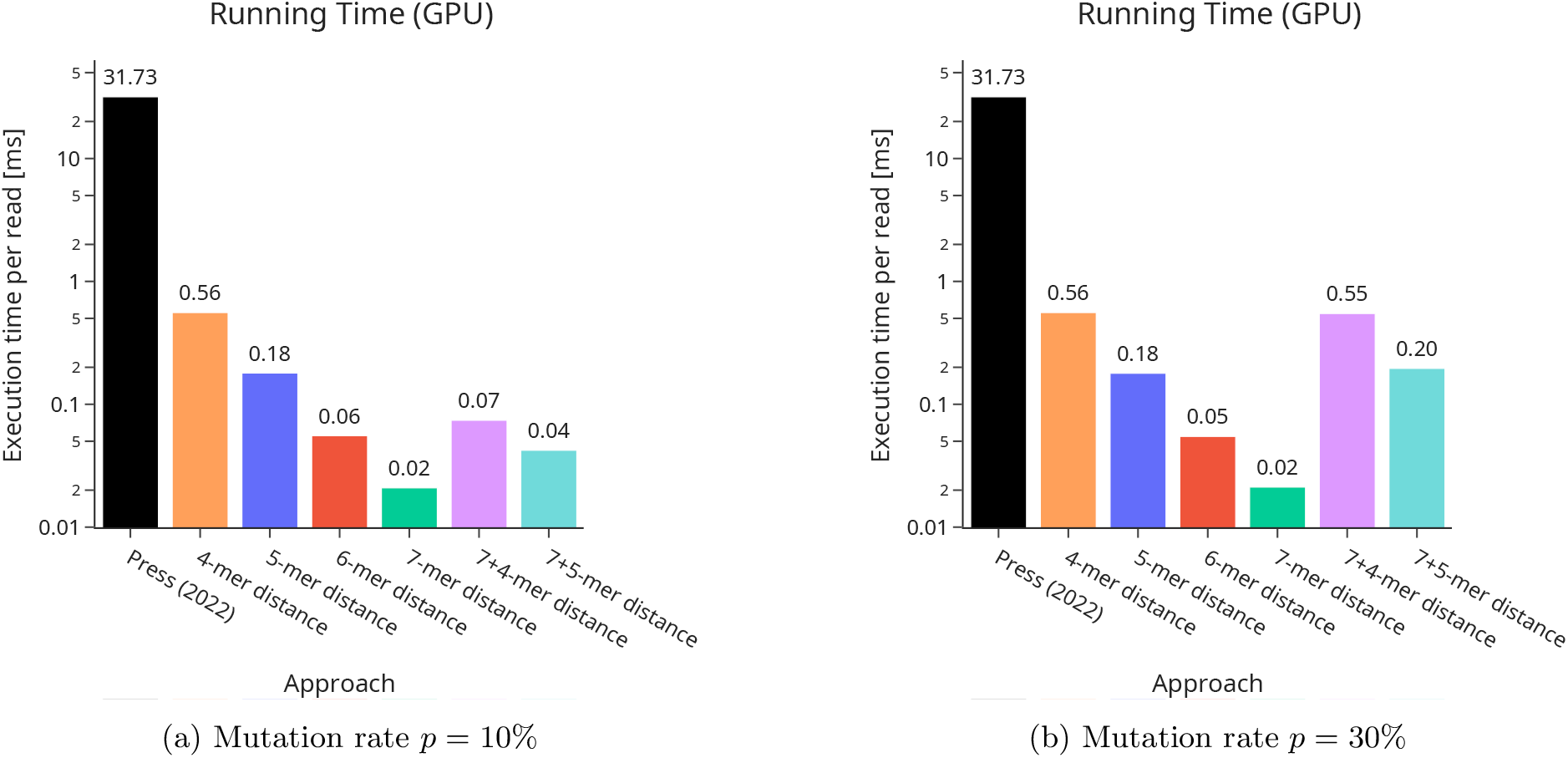
Execution times for *n* = 10^6^ barcodes on a GPU with all variants using the Levenshtein distance.

**Fig. 12:**
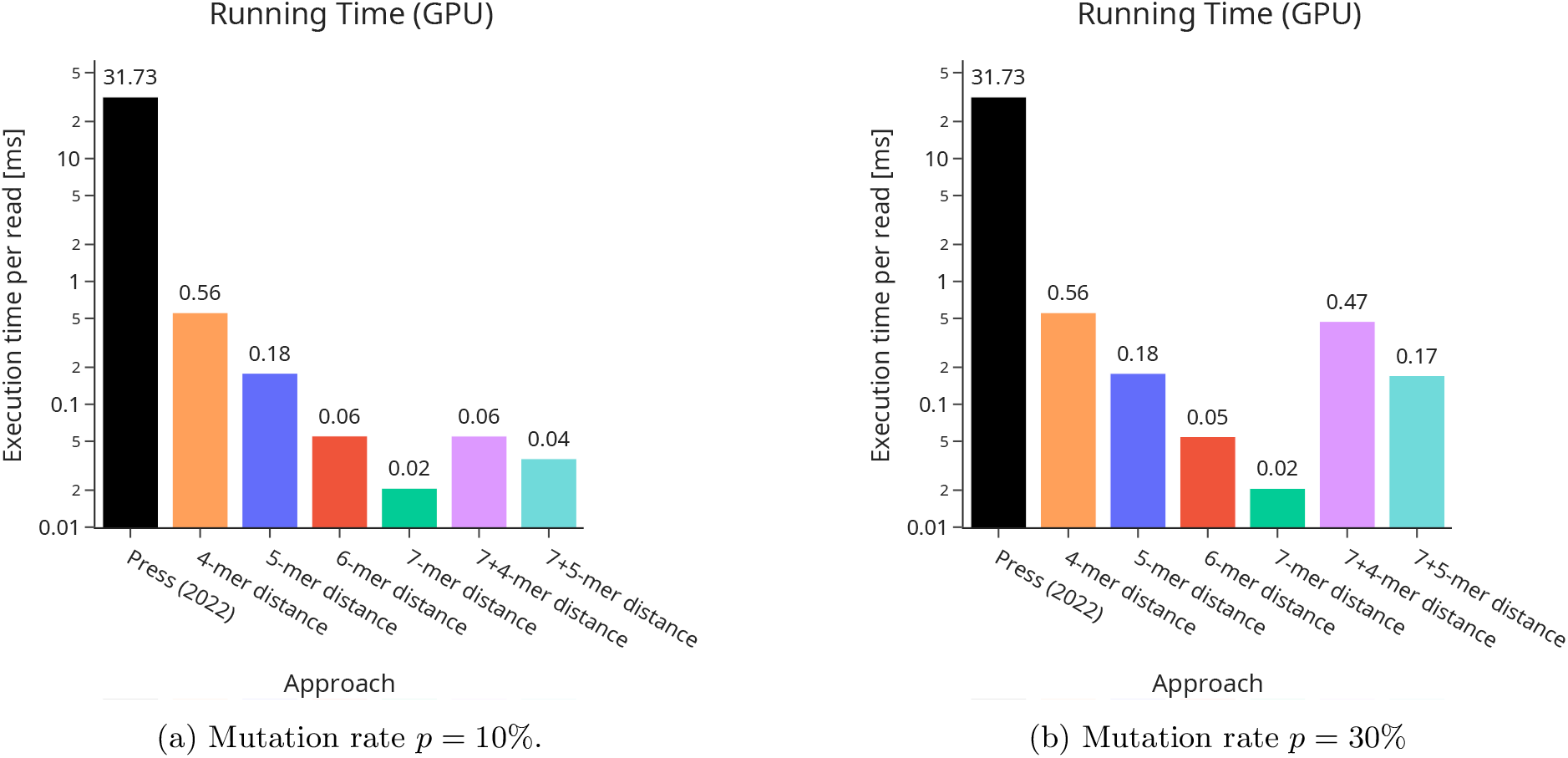
Execution times for *n* = 10^6^ barcodes on a GPU with the *k*-mer distance variants using the Sequence-Levenshtein distance.

### S3 Counting inner iterations for Press (2022)

Press (2022) [17] calculates two types of similarity measures (“triages”) between 3-mers (“trimers”) and later combines the candidate sets for each measure into a final ranking. For the triages, Press computes the Hamming distance between the 64-bit trimer occurrence vectors (“Hamming pop count”) and a cosine similarity measure between the positional encodings of the trimers for each read and barcode.

To calculate the Hamming pop count, one needs *n* inner iterations per read, one for each barcode, as shown in Algorithm 3. Furthermore, Press calculates a cosine similarity between the trimer position vectors of read and barcode (Algorithm 4). The position vectors have one entry for each possible trimer, totaling 64 entries. Thus, calculating the cosine similarity takes 64*n* inner iterations for each read. As Press computes four candidate sets using the cosine similarity with different parameters, the number of inner iterations for the triages sums up to *n* + 4 · 64*n* = 257*n* per read. Finally, the candidate sets of the triages are sorted and combined by multiplying the rank of the barcodes in each triage candidate set. The multiplied ranks serve as the final pseudo-distance, so the closest *c* barcodes make up the output candidate set of a read. Combining the triages requires another 5*n* inner iterations per read, as Algorithm 5 shows. This leaves us with 257*n* +5*n* = 262*n* inner iterations per read.

#### Algorithm 3 First triage: Hamming popcount

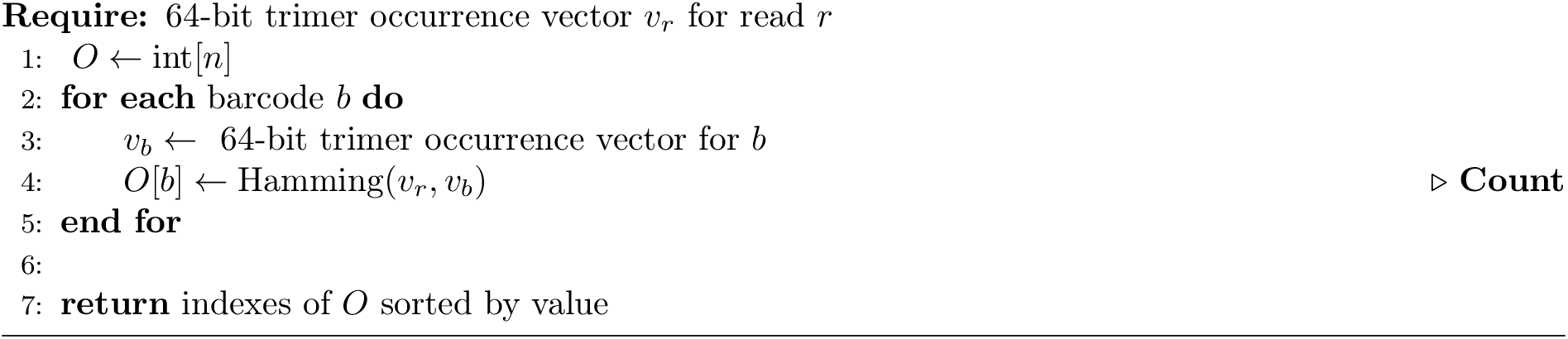

#### Algorithm 4 Second triage: Cosine similarity

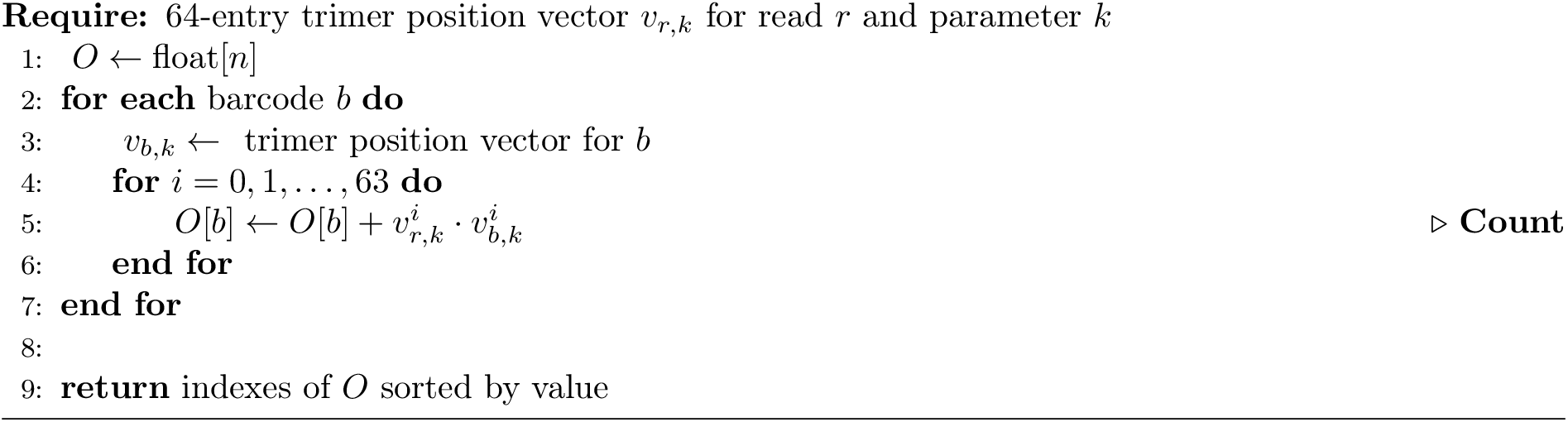

#### Algorithm 5 Combined triage

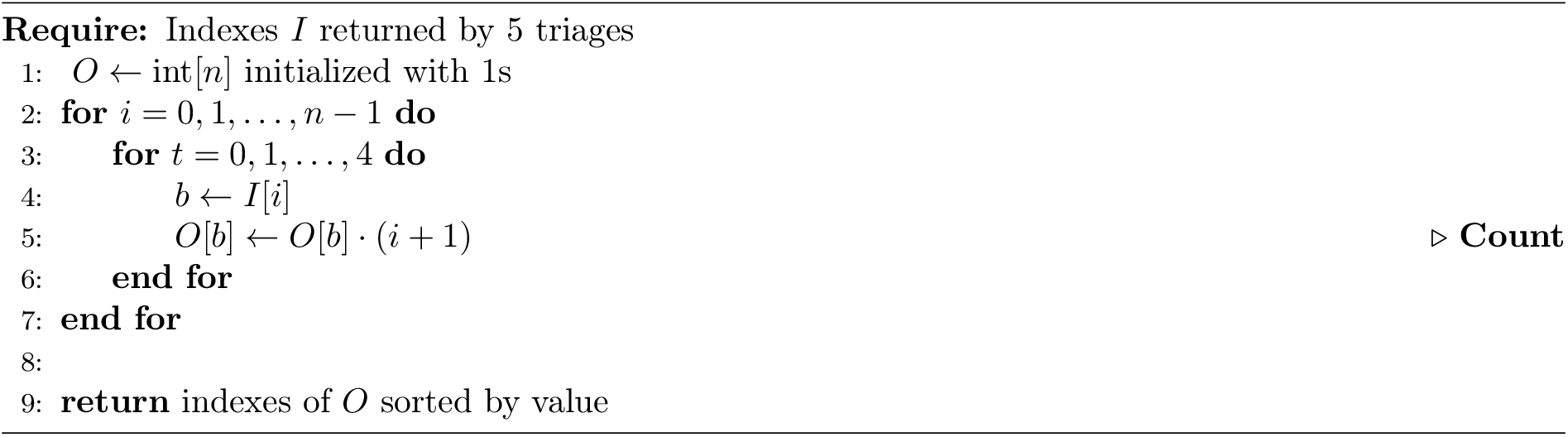

